# AlphaFold2 predicts interactions amidst confounding structural compatibility

**DOI:** 10.1101/2023.08.25.554771

**Authors:** Juliette Martin

## Abstract

Predicting physical interactions is one of the holy grails of computational biology, galvanized by rapid advancements in deep learning. AlphaFold2, although not developed with this goal, seems promising in this respect. Here, I test the prediction capability of AlphaFold2 on a very challenging data set, where proteins are structurally compatible, even when they do not interact. AlphaFold2 achieves high discrimination between interacting and non-interacting proteins, and the cases of misclassifications can either be rescued by revisiting the input sequences or can suggest false positives and negatives in the data set. Alphafold2 is thus not impaired by the compatibility between protein structures and has the potential to be applied at large scale.

## Introduction

Prediction of protein-protein interactions has profound implications to suggest functions for uncharacterized proteins, understand protein activity and regulation at the molecular level, and more generally, highlight protein functions in the context of global interactomes. Numerous computational methods have been developed to predict whether or not two proteins physically interact, based on their sequences and 3D structures, see the following references for review ^1–6^.

The formidable capability of AlphaFold2 (AF2) to predict protein 3D structures ^7^ has stimulated the creativity of the scientific community to evaluate what are the application range and limits of AF2 predictions ^8–20^ and extend the tool beyond its initial prediction task ^18, 21–38^. Prediction of protein-protein complex structures, a task traditionally addressed by protein-protein docking, has rapidly been tackled by modification in the input for the AF2 monomer pipeline ^24^. A specific model for protein-protein complexes is now available, with breakthrough prediction results ^39^. Note that in this case, predictions are made with the prior knowledge that the proteins physically interact.

Logically then, the capability of AlphaFold2 to predict interaction between proteins has recently been explored, by using the predicted quality of modeled interfaces as prediction criterion, with very encouraging discrimination capability ^28, 35, 40, 41^.

In this short article, I challenge AF2 on a particular, presumably difficult data set in which non-interacting protein pairs are special cases, in which the two (non-interacting) proteins are structurally similar to available experimental complexes ^42^. This feature should challenge AF2, since the proteins are compatible in terms of structures. Using the iPTM score of AF2, I found that AF2 is very accurate at discriminating interacting from non-interacting pairs, even in this challenging context, attending an AUC value of 0.93. Interestingly, model recycling did not improve the discriminative power. The analysis of the few misclassified cases provides suggestions to further improve the discrimination and how to use AF2 for pair screening.

## Material and Methods

### Data set

Protein pairs from *S cerevisiae* are taken from our previous study ^42^, details about these datas set can be bound in ^42^ and briefly summarized below.

### Interacting protein pairs

The initial data set of interacting protein pairs was extracted from three sources: high confidence physical interactions from BioGrid ^43^, direct interactions from the KUPS resource ^44^, and high confidence physical interactions detected by yeast-two-hybrid from Ito et al ^45^.

### Non-interacting protein pairs

The initial negative data set was extracted from three sources : the negative data set used by Yu et al which are simply sampled from pairs without experimental evidence of interaction ^46^, pairs of protein from the KUPS resource ^44^, which have no evidence of interaction and also distant GO annotations, and the negative data set built by Trabuco et al ^47^ from the Ito data set, where the yeast-two-hybrid data set is used to select proteins pairs without interaction but correctly detected in the experiment .

In our previous study, those pairs were compared with known structures. We had screened homology models of S. *cerevisiae* proteins against a non-redundant database of experimentally known dimers and we had selected pairs where the monomers structurally matched with the experimental dimers (TM score >0.8), and, once superimposed on those dimers, could form an interface of reasonable size (> 20 residues) and without extensive clashes (less than 3 between Cαs). Using these criteria resulted in a data set of 22 non-interacting and 222 interacting proteins. In this work, I use the data set of 22 non-interacting proteins and a random sample of 22 interacting pairs, see Table S1.

### AlphaFold2 Models

AlphaFold2 predictions are computed using LocalColabFold (https://github.com/YoshitakaMo/localcolabfold), a local installation of ColabFold ^39, 48^. ColabFold replaces the time-consuming step of multiple sequence alignment (MSA) creation by an ultra-fast step with MMseqs2 ^49^. No templates are used; models are not minimized; I tested both with and without model recycling, and different modes of sequence pairing for the MSA: unpaired+paired (default), paired only, and unpaired only.

The resulting 5 models are ranked according to the ipTM computed by AF2 and this score is used as a predictor of protein interaction.

### Assessing classification performance

The separation between scores of interacting and non-interacting pairs is measured by the AUC value when using the score to predict interaction. This is done for the AF2 ipTM score and the pDockQ, recently introduced by Bryant et al ^40^, which is a derived from the plDDT scores of interface residues. Statistical significance between AUC values is assessed using the non-parametric DeLong’s test ^50^ implemented in the pROC package ^51^.

The classification performance is measured by the accuracy, i.e., percentage of correctly classified pairs. Statistical difference between accuracies is assessed using the MacNemar test ^52^.

## Results

Although AF2 has recently been used to discriminate interacting from non-interacting pairs with promising results ^28, 35, 40, 41^, it is always worthy of pushing the system to the limits to better know its applicability range. Here, I propose to further test AF2 prediction capability in extreme conditions. I submitted to AF2 prediction a particularly challenging data set from a previous study ^42^. In this data set, all the pairs are supported by structural data:, interacting but also non-interacting pairs are compatible in shape, as assessed by their high similarity to experimental dimers, as explained in the Methods section.

The results of AF2 prediction on this challenging data set are impressive, attaining AUC value in the 0.86-0.93 range, see Figure 1, and accuracy around 80-86%. These results are comparable to the recent result of Bryant et al who obtained an AUC equal to 0.87 using the monomer AF2 pipeline on E. *coli* proteins ^41^. Detailed results are shown in Tables S2 and S3 for each AF2 model, with and without recycling, and different MSA pairing modes. Unfortunately, the limited size of the data set does not allow to statistically differentiate those different settings: the predictions are uniformly good. AF2 ipTM score outperforms the pdockQ score in terms of separation (Delong’s test p-value=0.011, see Table S3). Considering the predictions obtained without recycling, default MSA pairing and the best model out of five according to the ipTM score, 6 pairs are misclassified, corresponding to an accuracy equal to 86%. The misclassified cases are shown in Figures 2 and 3 and discussed below.

**Figure 1.**
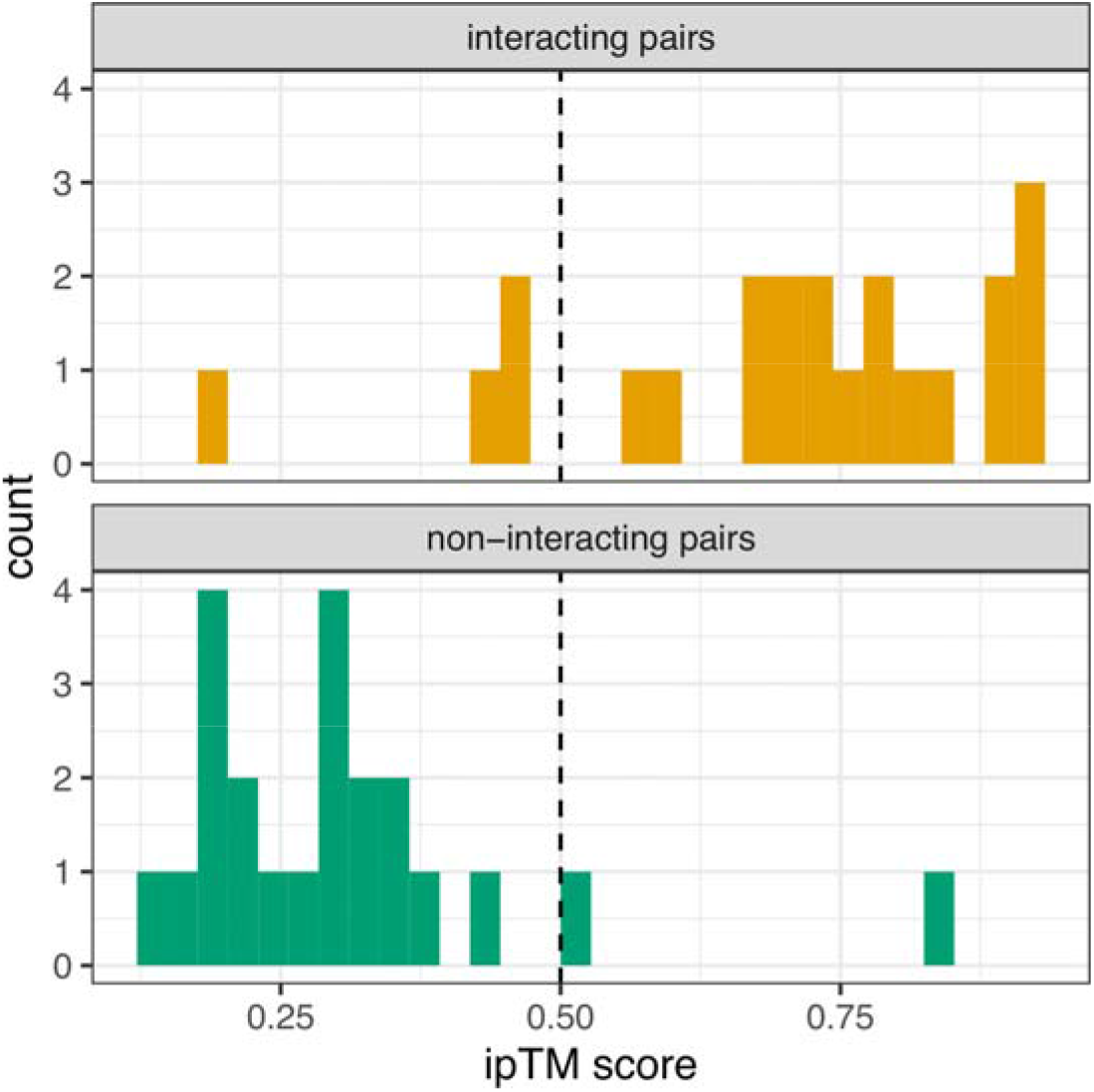
iPTM score distribution for interacting and non-interacting pairs. The vertical dashed line indicates the prediction cutoff (0.5).

**Figure 2.**
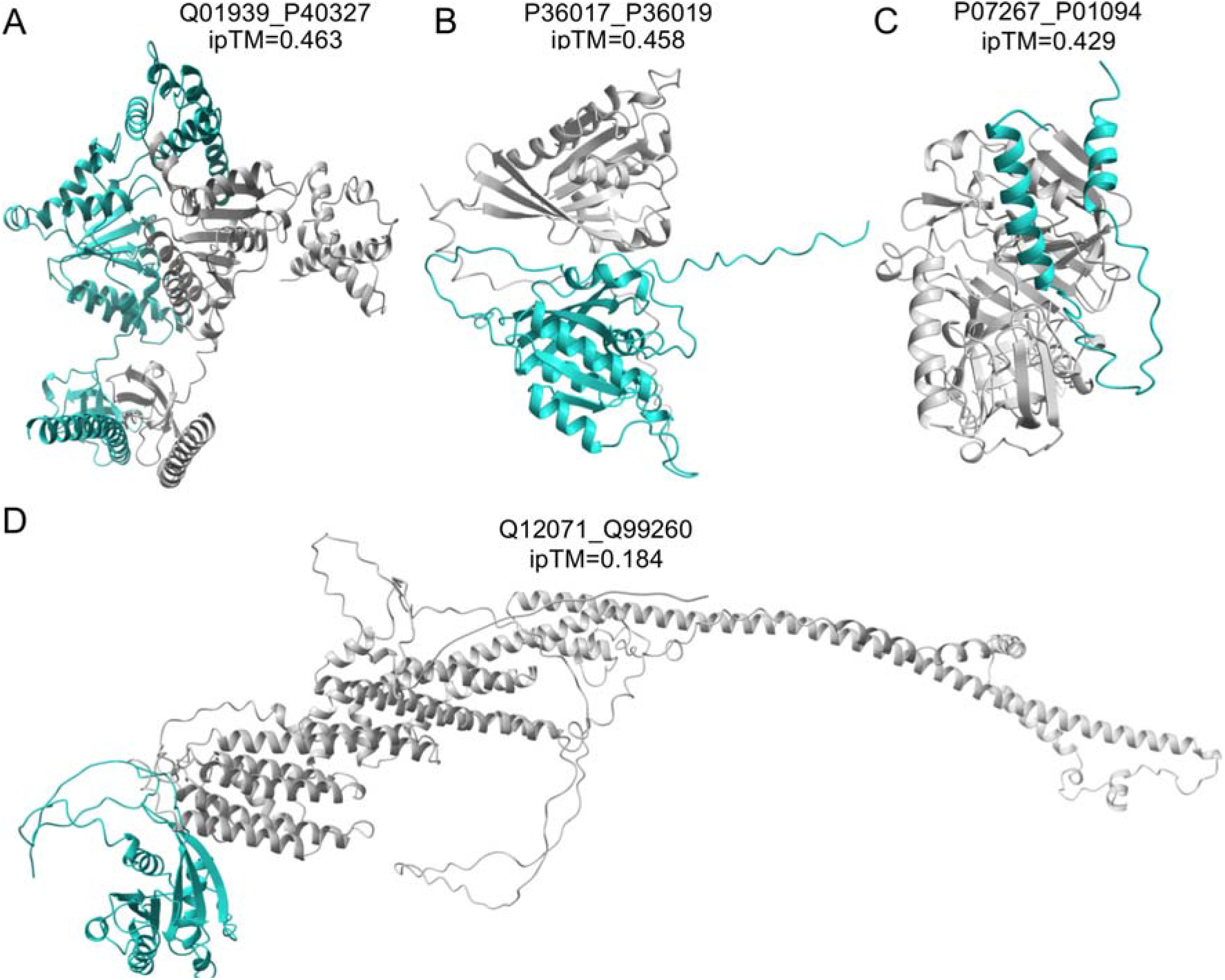
AF2 models of interacting pairs misclassified as non-interacting (ipTM<0.5). A: pair formed by the subunits 8 (Q01939, in blue) and 4 (P40327, in grey) of the 26s proteasome. B: pair formed by the vacuolar protein sorting-associated protein 21 (P36017 in grey) and the GTP-binding protein YPT53 (P36019, in blue). C: pair formed by saccharopepsin (P07267, in grey) and its inhibitor (P01094, in blue). D: pair formed by the vacuolar protein sorting-associated protein 54 (Q12071, in grey) and the GTP-binding protein YPT6 (Q99260, in blue).

**Figure 3.**
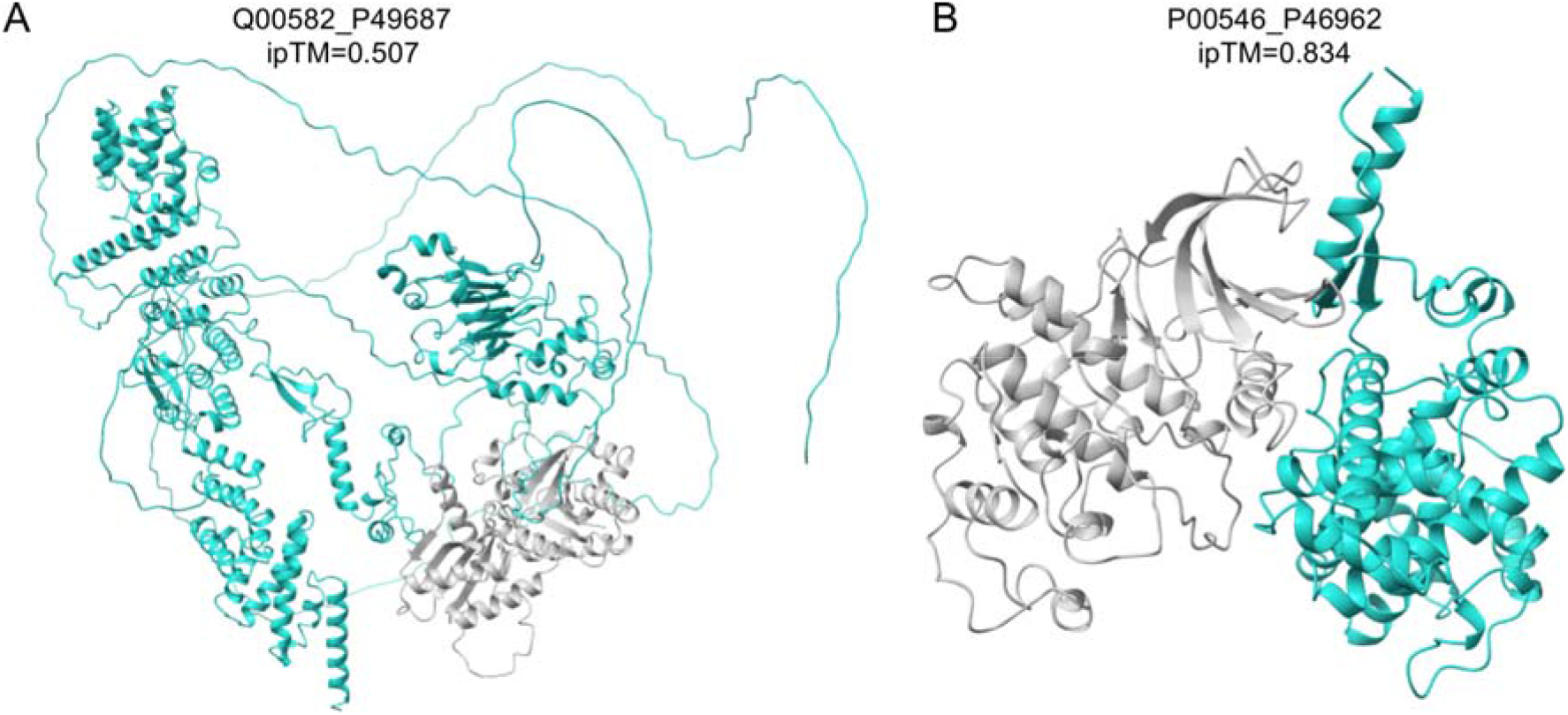
AF2 models of non-interacting pair misclassified as interacting (ipTM>0.5). A: pair formed by the GTP-binding protein GTR1 (Q00582, in grey) and the nucleoporin NUP145 (P49687, in blue). B: pair formed by cyclin-dependent kinase 1 (P00546, in grey) and the CTD kinase subunit beta (P46962, in blue)

Four interacting pairs are incorrectly classified as non-interacting (ipTM score <0.5). The first case is the dimer formed by the regulatory subunit 8 homolog and subunit 4 homolog (Uniprot ids Q01939 and P40327) of the 26S proteasome, which is an assembly of 47 protein chains. The dimer obtains an ipTM score equal to 0.463, close to the interaction cutoff, see Figure 2A. The comparison of the AF2 model with the experimental structure of the 26S proteasome (PDB id 3JCO) reveals that the AF2 model is structurally similar to the dimer formed by the subunits 7 and 4, meaning that AF2 wrongly placed the subunit 8 in place of the subunit 7, see Figure S1. This is possible because subunits 8 and 7 are structurally similar (TM score = 0.6 between experimental structures). So, in this case, AF2 prediction was confused by the presence of another interacting chain with similar structure.

The pair formed by the vacuolar protein sorting-associated protein 21 and the GTP-binding protein YPT53 (Uniprot ids P36017 and P36019) obtains an ipTM score equal to 0.458, close to the prediction cutoff. Furthermore, the PAE matrix displays low values between the two chains, suggesting a good confidence in the relative orientation of the two protein chains. The AF2 model, shown in Figure 2B, has an interface involving the disorder C-tail of the GTP protein, which could explain the low score. Interestingly, the model obtained with recycling has a good ipTM score (0.616) with a similar configuration but without the disordered part at the interface (see Figure S2). However, model recycling significantly increases the computation time, which is a limiting factor in the perspective of pair screening. Alternatively to model recycling, the prediction can be run with the disordered part chopped from the sequence, which produces a model similar to the recycled one, with an ipTM score equal to 0.63 (see Figure S2). So, in that case, it is possible to ‘rescue’ the prediction by chopping the sequence.

The pair formed by the saccharopepsin and its inhibitor (Uniprot ids P07267 and P01094) obtains an ipTM score equal to 0.429, see Figure 2C. The full-length sequence of the saccharopepsin contains an N-terminal propeptide of 75 residues that is cleaved upon activation of the enzyme ^53^. The comparison of the AF2 model with the experimental structure (PDB id 3COJ) reveals that the binding cleft where the inhibitor is supposed to bind is occluded by the N-terminal region of the enzyme corresponding to the propeptide. Re-running the prediction after chopping the propeptide sequence results in a model with a good ipTM score equal to 0.63 and in good agreement with the experimental structure (see Figure S3). So, in this case also, it is possible to rescue the prediction with appropriate sequence chopping.

The pair formed by the vacuolar protein sorting-associated protein 54 and the GTP-binding protein YPT6 (Uniprot ids Q12071 and Q99260) obtains a very low ipTM score equal to 0.184, see Figure 2D. The 5 models are drastically different from each other, with even drastic situations where the proteins are not in contact in two of the models (see Figure S4). There is evidence of physical interaction between these proteins, as detected by affinity purification. However, there is no evidence of direct physical interaction by two-hybrid assay. So there is the possibility that this pair is in fact a false positive case.

Two pairs of non-interacting proteins are incorrectly classified as interacting. The pair between GTP-binding protein GTR1 and nucleoporin NUP145, a component of the nuclear pore complex (Uniprot id Q00582 and P49687) obtains a border line ipTM score equal to 0.507, see Figure 1F. The examination of the AF2 model reveals that the nucleoporin has a disordered N-terminal region that mediates the interaction with GTR1. After chopping the disordered parts of the nucleoporin, the best model obtains an ipTM score equal to 0.23. So in this case, the initial ipTM score was meaningless because of the disordered parts, and the interaction can be excluded by adequate sequence chopping.

The pair formed by the cyclin-dependent kinase 1 and the CTD kinase subunit beta (Uniprot id P00546 and P46962) achieves a very high ipTM score equal to 0.834, see Figure 1G. In this case, all models have high ipTM scores (>0.8) and are structurally similar (data not shown). Although there is no evidence of direct interaction between these two proteins, they are reported as interacting in the STRING resource ^54^, having, among other things, a direct interaction between homologs in Drosophila measured by yeast-two-hybrid assay ^55^. This suggests that this case could be a false negative pair.

In summary, out of 6 misclassified cases, three could be corrected by sequence chopping, two can be questioned as false positive/negative, and one highlights a phenomenon of confusion between chains in interfaces in a macromolecular assembly.

## Conclusion

A reduced but challenging data set was submitted to AF2 in order to discriminate interacting from non-interacting pairs, resulting in very high prediction accuracy. Several misclassified cases could be rescued by appropriate sequence chopping, and some others are suggestive of incorrect annotations (false positive or false negative). A potential limitation of AF2 was observed in a case where several protein chains with structural similarity form a supra-molecular assembly. The fact that no recycling is required opens the possibility to apply this procedure at large scale. To conclude, AF2 seems a promising technology for predicting protein-protein interactions, even capable of discriminating interacting from non-interacting pairs in presence of confounding structural compatibility.

