## Supplementary material for "AlphaFold2 predicts interactions amidst confounding structural compatibility": Tables S2 to S4, Figures S1 to S4

Juliette Martin

**Table S2**- AUC values obtained with AF2 confidence score used as a predictor. The differences between AUC values are not statistically significant between different MSA pairing and recycling modes using the best network (last line), or between different networks in each MSA/recycling setting (each column).

| | MSA pairing mode | paired+unpaired | | paired only | unpaired only |
|---|---|---|---|---|---|
| | recycling mode | With recycling | No recycling | No recycling | No recycling |
| AlphaFold network | 1 | 0.9215 | 0.8905 | 0.7893 | 0.9081 |
| | 2 | 0.9298 | 0.8698 | 0.8202 | 0.8812 |
| | 3 | 0.9132 | 0.906 | 0.7593 | 0.8244 |
| | 4 | 0.8967 | 0.8709 | 0.7665 | 0.9514 |
| | 5 | 0.8843 | 0.9112 | 0.874 | 0.9339 |
| | Best | 0.8636 | 0.9256 | 0.8595 | 0.9143 |

**Table S3**- Accuracy values obtained with AF2 confidence score as a predictor, and a cutoff equal to 0.5. The differences between accuracy values are not statistically significant between each other according to the McNemar test.

| | MSA pairing mode | paired+unpaired | | paired only | unpaired only |
|---|---|---|---|---|---|
| | recycling mode | With recycling | No recycling | No recycling | No recycling |
| AlphaFold network | Best | 80% | 86% | 84% | 84% |

**Table S4**- AUC values obtained with pDockQ score used as a predictor. The difference between AUC values (without recycling, paired+unpaired MSAs) is significant between AF2 confidence scores and pDockQ scores (0.9256 versus 0.6684, Delong's test p-value=0.011).

| | MSA pairing mode | paired+unpaired | | paired only | unpaired only |
|---|---|---|---|---|---|
| | recycling mode | With recycling | No recycling | No recycling | No recycling |
| AlphaFold network | 1 | 0.8502 | 0.7913 | 0.8698 | 0.8554 |
| | 2 | 0.8461 | 0.7304 | 0.8326 | 0.8347 |
| | 3 | 0.7727 | 0.7149 | 0.7831 | 0.7727 |
| | 4 | 0.7893 | 0.7727 | 0.7944 | 0.7386 |
| | 5 | 0.7851 | 0.8161 | 0.7996 | 0.8140 |
| | Best | 0.7758 | 0.6684 | 0.78 | 0.7149 |

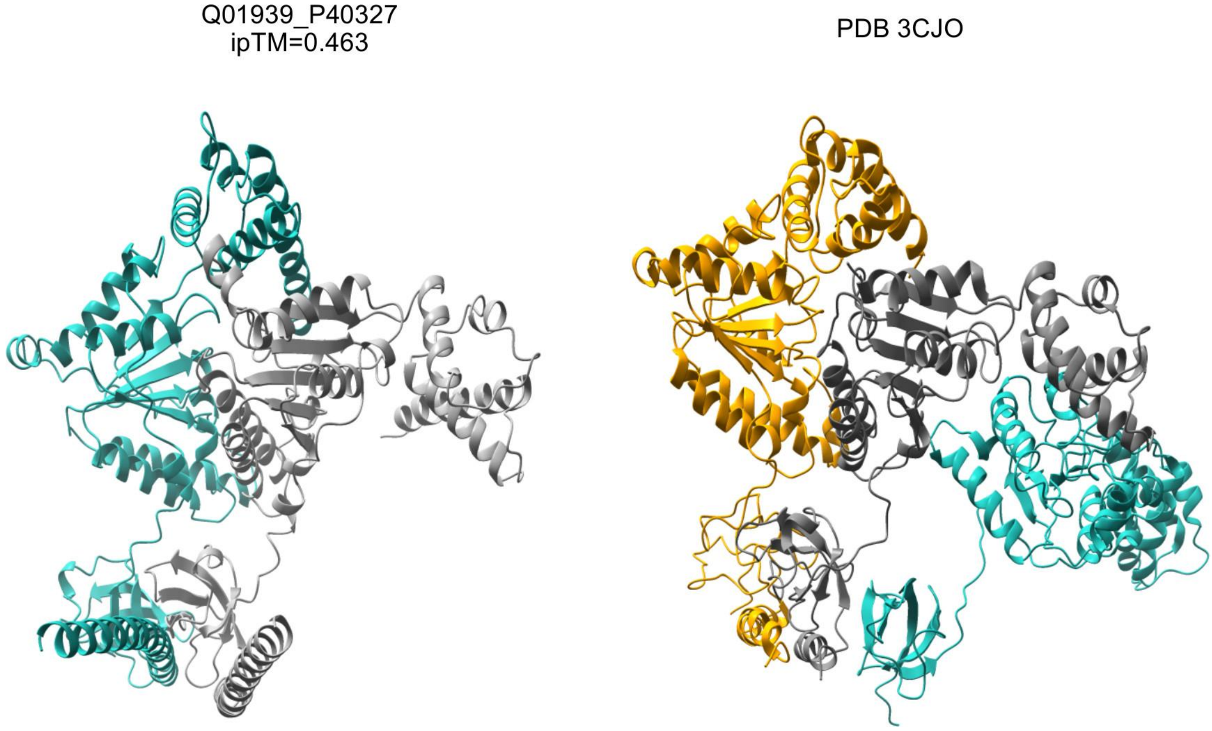


Figure S1. Left: the AF2 model of the misclassified interacting pair formed by the subunits 8 (Q01939, in blue) and 4 (P40327, in gray) of the 26S proteasome. Right: a subset of the experimental structure of the 26S proteasome, with subunit 8 in blue, subunit 4 in gray, and subunit 7 in orange.

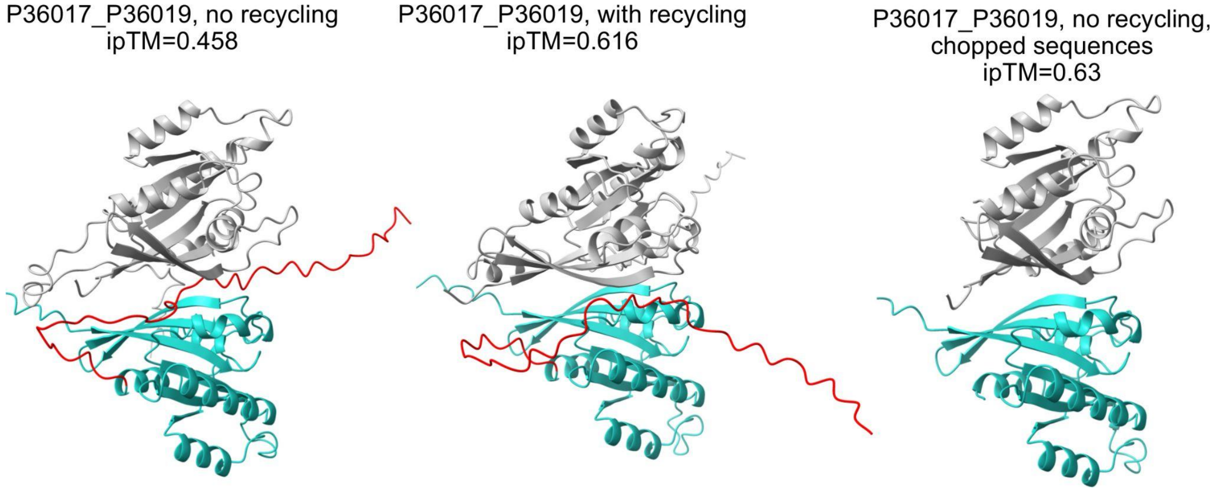


Figure S2. AF2 models of the misclassified interacting pair formed by the vacuolar protein sorting-associated protein 21 (P36017, in gray) and the GTP-binding protein YPT53 (P36019 in blue, with the C-terminal disordered part in red).

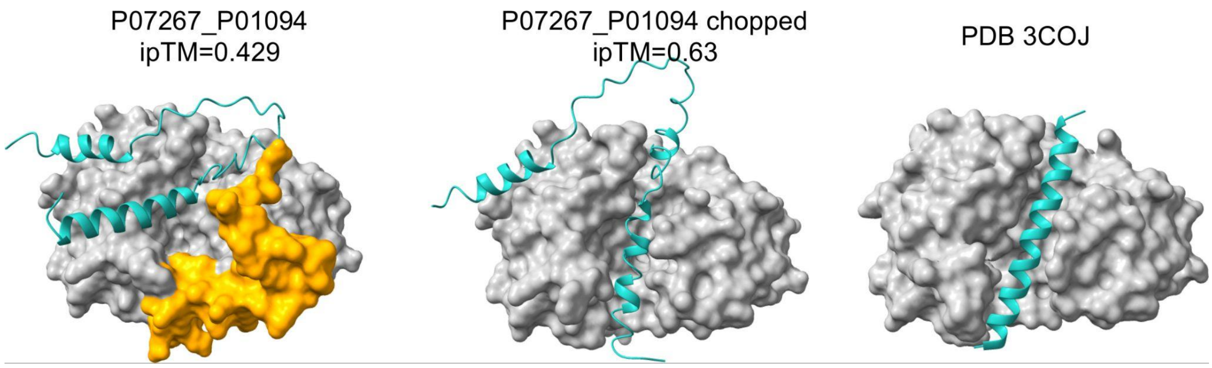


Figure S3. AF2 models of the misclassified interacting pair formed by the saccharopepsin (P07267, in gray, and the propeptide in orange) and its inhibitor (P01094, in blue), together with the experimental structure of the complex.

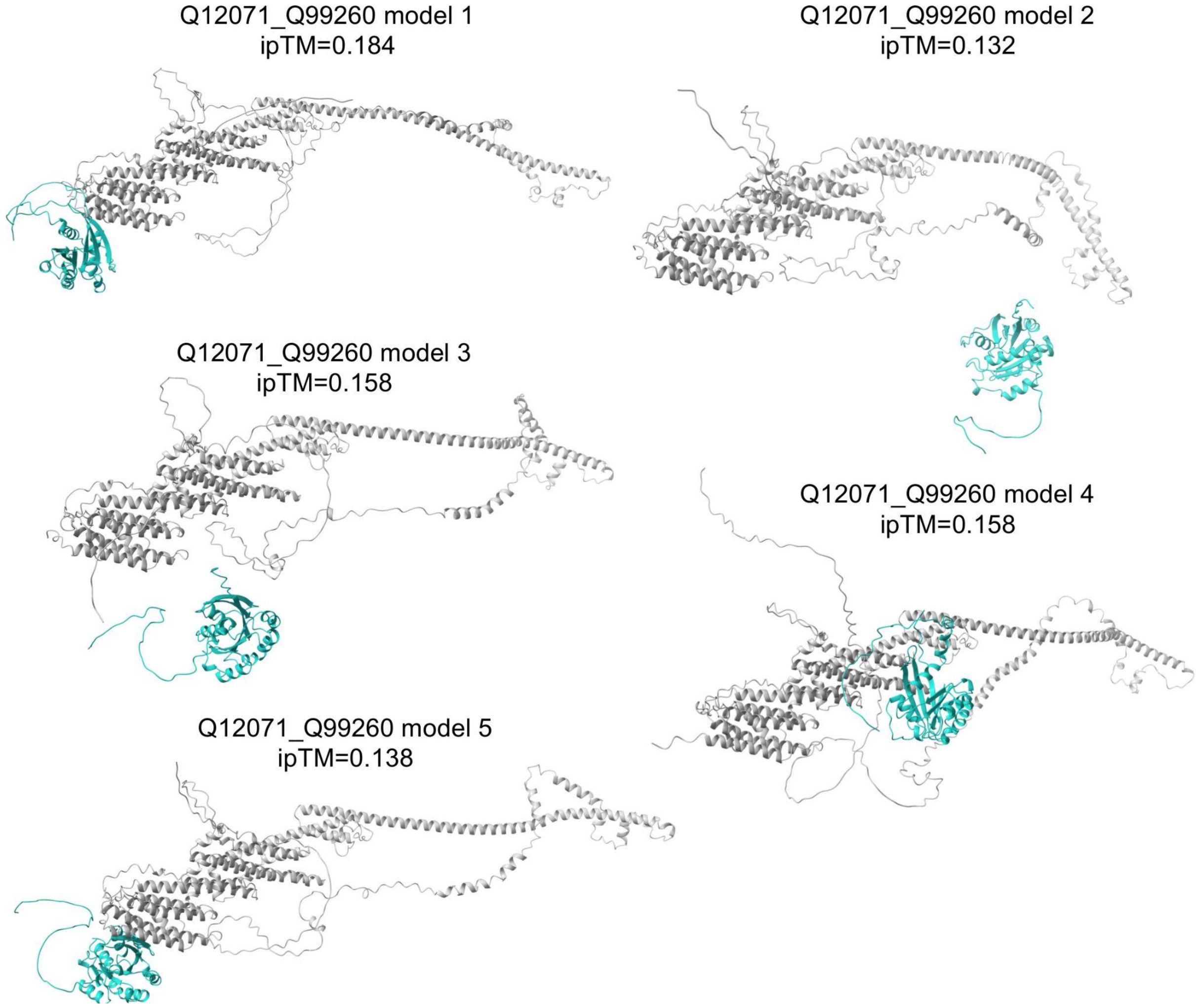


Figure S4. AF2 models for the misclassified interacting pair formed by the vacuolar protein sorting-associated protein 54 (Q12071, in gray) and the GTP-binding protein YPT6 (Q99260, in blue). Proteins do not form contacts in model 2 and model 5.
